# Integrative network-based construction of ecologically relevant computational adverse outcome pathways for organic mercury-induced toxicity

**DOI:** 10.1101/2025.10.25.684520

**Authors:** Shreyes Rajan Madgaonkar, Nikhil Chivukula, Vasavi Garisetti, Shambanagouda Rudragouda Marigoudar, Krishna Venkatarama Sharma, Areejit Samal

## Abstract

Adverse outcome pathways (AOPs) describe mechanisms of toxicity by connecting molecular events with outcomes at higher levels of biological organization. Computational AOPs (cAOPs), constructed using existing toxicological data, can accelerate the early stages of AOP development, which is often a time-consuming and resource-intensive process. In this study, an integrative network-based framework was developed to construct cAOPs, with a particular focus on elucidating the toxicity of organic mercury in fish. First, 124 organic mercury compounds, corresponding fish-specific toxicity endpoints, and proteins were curated from CTD and ECOTOX. Next, molecular docking investigations were performed to determine novel molecular interactions with 16 zebrafish protein orthologs. Subsequently, the toxicity endpoints, including the identified molecular interactions, were standardized and harmonized using data from AOP-Wiki, Gene Ontology, and MeSH. These endpoints were integrated with event relationship information from AOP-Wiki and published literature to construct an organic mercury-associated toxicity network comprising 197 nodes and 243 edges. Thereafter, node- and edge-level filtration criteria were designed based on AOP definitions and utilized to identify biologically relevant pathways within the toxicity network. Further, these pathways were ranked based on their novelty with respect to existing AOPs within AOP-Wiki. Finally, based on extensive evidence from published literature, four top-ranked novel pathways that describe binding with glutathione peroxidase or alterations in metallothionein levels leading to neurologic manifestations or dysbiosis in fish were proposed as cAOPs. Overall, this study presents an integrative network-based framework for constructing cAOPs applicable to diverse contaminants and species, supporting New Approach Methodologies for toxicological risk assessment.

## 1. Introduction

Adverse Outcome Pathway (AOP) is a stressor-agnostic framework that organizes mechanistic toxicological information by linking biological events across multiple levels of organization, beginning at the molecular level and progressing to adverse outcomes at higher levels (Ankley et al., 2010; Villeneuve et al., 2014a). AOPs aim to reduce reliance on animal testing through the development of *in vitro* assays and *in silico* models, thereby advancing new approach methodologies (NAMs) in toxicity testing (Leist et al., 2017; Tollefsen et al., 2014). Combining these strategies facilitates integrated approaches to testing and assessment (IATA) for improved hazard identification, chemical prioritization, and risk assessment (Tollefsen et al., 2014). AOP-Wiki (https://www.aopwiki.org) serves as the largest global repository of AOPs and is maintained by the Organisation for Economic Cooperation and Development (OECD). AOPs are submitted to this repository by researchers worldwide, where they undergo extensive scientific review before receiving formal endorsement status. These AOPs have been utilized to investigate complex biological mechanisms underlying toxicities caused by diverse chemical stressors (Aguayo-Orozco et al., 2019; Jeong and Choi, 2019; Kim et al., 2024; Sahoo et al., 2024b, 2024a, 2024c). Overall, AOPs provide a systematic approach to integrate mechanistic knowledge and support evidence-based risk assessment strategies.

However, existing AOPs cover only a small portion of toxicologically relevant biological space (Bell et al., 2016), with ecotoxicologically relevant pathways being particularly underrepresented (Sahoo et al., 2024c). This limitation stems from the time- and resource-intensive process of AOP development that involves extensive experimental investigation (Leist et al., 2017) and demands a comprehensive literature review (Bell et al., 2016) to enable in-depth analysis and ensure the scientific validity of the proposed pathway. The subsequent formal evaluation and endorsement process further extends the timeline (Svingen et al., 2021). To accelerate early=stage AOP development, computational approaches have been proposed that extract toxicity data from publicly available resources and infer associations to produce adverse outcome-specific computational AOPs (cAOPs) (Baker et al., 2020; Bell et al., 2016; Bozic et al., 2023; Fan et al., 2024; Jeong et al., 2019; Jeong and Choi, 2020; Oki and Edwards, 2016; Wang et al., 2022; Zhang et al., 2022). Although AOP-Wiki has not yet been widely integrated into computational approaches, its structured content provides a valuable framework for organizing pathways and highlighting their novelty. However, due to inherent redundancies and inconsistencies, AOP-Wiki does not fully align with FAIR (Findable, Accessible, Interoperable, and Re-usable) data sharing principles (Wilkinson et al., 2016), making its integration with external resources challenging. Therefore, acceleration of the AOP development process necessitates improved computational strategies that can effectively manage redundancies and enable systematic integration of AOP-Wiki data with publicly available resources.

Mercury is recognized as a highly toxic environmental contaminant globally (Balali-Mood et al., 2021; Clarkson and Magos, 2006; Driscoll et al., 2013; Wu et al., 2024) and is ranked third on the Agency for Toxic Substances and Disease Registry (ATSDR) Substance Priority List (SPL) due to its widespread occurrence, severe toxicity, and significant potential for human exposure (https://www.atsdr.cdc.gov/programs/substance-priority-list.html). It persists in the environment and can be transformed into organic forms, such as methylmercury, that can readily bioaccumulate and biomagnify in aquatic food webs (Budnik and Casteleyn, 2019; Jeong et al., 2024; Lavoie et al., 2013; Obrist et al., 2018). Despite extensive regulatory efforts, organic forms of mercury continue to pose substantial ecological and human health risks owing to their high persistence and toxicity within aquatic ecosystems (Jeong et al., 2024; Kayani and Mohammed, 2025; Wu et al., 2024). It also endangers the health and livelihoods of human populations relying on marine and freshwater resources, an essential pillar of the blue economy (IRP, 2021). A prominent example of the severe consequences of organic mercury exposure is Minamata disease, first identified in Japan during the 1950s, where widespread methylmercury poisoning resulted in severe neurological disorders and fatalities among local communities consuming contaminated seafood (Yokoyama, 2018). Moreover, organic mercury-associated toxicity mechanisms have been explored through the AOP framework in humans. For instance, methylmercuric chloride is listed as one of the prototypical stressors for the endorsed AOP:17 (https://aopwiki.org/aops/17), titled ‘Binding of electrophilic chemicals to SH(thiol)-group of proteins and /or to seleno-proteins involved in protection against oxidative stress during brain development leads to impairment of learning and memory’, within AOP-Wiki. Similarly, Li *et al*. constructed an AOP to investigate methylmercury-induced congenital diseases and developmental failure, through experimental studies on human embryonic stem cells (Li et al., 2023). While these efforts consolidate organic mercury-induced toxicity data in humans, comparable ecotoxicological data for aquatic species remain largely unorganized and underexplored. Addressing this gap is vital as the bioaccumulation of organic mercury within aquatic food webs and its subsequent biomagnification threatens not only ecosystem integrity (de Almeida Rodrigues et al., 2019; UN Environment, 2019), but also the safety of human populations relying on these resources for sustenance (M.-L. Li et al., 2024; UN Environment, 2019), which makes organic mercury an ideal stressor for developing cAOPs.

In this study, an integrative network-based approach was developed to construct ecotoxicologically-relevant cAOP for organic mercury-induced toxicity. First, toxicological databases such as CTD and ECOTOX were utilized to identify chemicals along with their associated toxicity endpoints and protein targets. Next, zebrafish protein orthologs were identified, and molecular docking investigations were performed to determine novel molecular interactions. The toxicity endpoints and these events were then standardized to initialize a toxicity network. The AOP-Wiki and published evidence were used to identify the edges in the network. Subsequently, functionally similar events were harmonized to mitigate redundancies, resulting in the construction of an organic mercury-specific toxicity network. Thereafter, node- and edge-level criteria were designed based on AOP definitions and utilized to identify biologically relevant paths within this toxicity network. Finally, these paths were compared against existing AOP information to assess their novelty, and the most novel paths were supported with evidence from published literature and proposed as cAOPs. Overall, this study presents a novel network-centric methodology based on AOP development guidelines that integrates ecotoxicity data, predicted molecular interactions, and existing AOP information to facilitate the construction of cAOPs for fish, with organic mercury serving as a prototypical stressor. Moreover, this approach can be extended to other contaminants and species, providing a framework to investigate their toxicological effects and support their mitigation strategies.

## 2. Materials and Methods

### 2.1. Compilation and curation of organic mercury compounds

Organic forms of mercury are legacy pollutants that pose significant health risks due to their high toxicity and ability to bioaccumulate (Kayani and Mohammed, 2025; Wu et al., 2024). While the effects on human health are well-documented, a critical knowledge gap exists regarding the impact of organic mercury on aquatic organisms, which is crucial as they serve as the primary entry point into food webs, threatening ecosystem integrity and human safety (UN Environment, 2019; Zheng et al., 2019). To bridge this gap, this study presents an integrative network-based framework that compiles existing toxicology information into a toxicity network and leverages the toxicity network to construct computational Adverse Outcome Pathways (cAOPs) elucidating the ecotoxicity of organic mercury in fish (Figure 1). As a first step, organic mercury compounds with documented toxicity endpoints were compiled from two databases, the Comparative Toxicogenomics Database (CTD) (https://ctdbase.org/) and the ECOTOX knowledgebase (https://cfpub.epa.gov/ecotox/).

**Figure 1:**
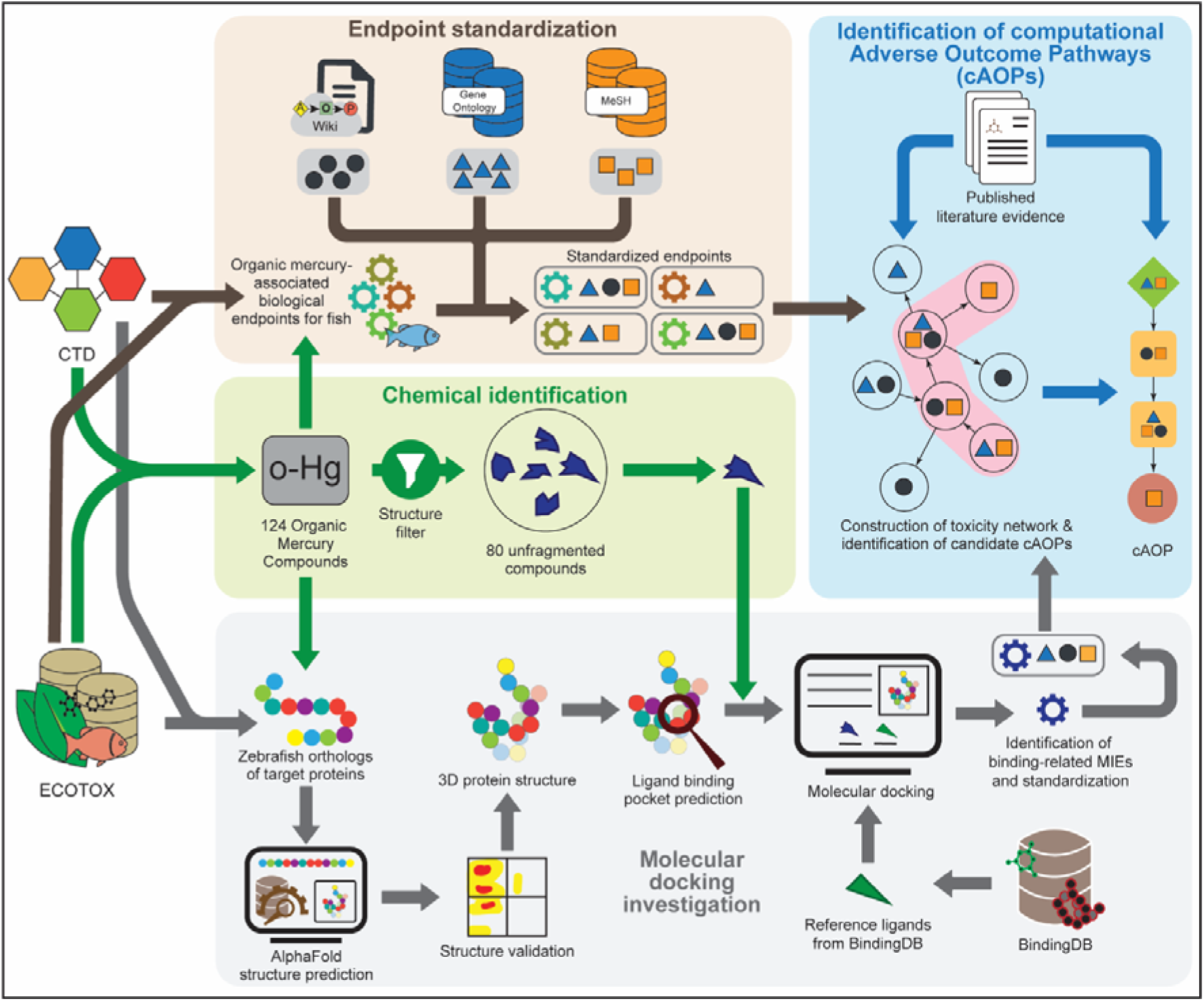
Summary of workflow followed to identify organic mercury compounds, followed by molecular docking investigation, standardization of organic mercury-associated biological endpoints in fish, and identification of computational Adverse Outcome Pathways (cAOPs).

CTD provides information on genes and/or proteins, phenotypes, and diseases associated with exposure to a chemical in different species. ECOTOX is a database built by the United States Environmental Protection Agency (US EPA) that curates information from ecotoxicity studies, including test species, toxicity endpoints, and their concentration values, among other details. First, all chemicals under class ‘Organomercury Compounds’ (https://ctdbase.org/detail.go?type=chem&acc=D009941#treeD02.691.750) in CTD, and under category ‘Mercury’ in ECOTOX, were extracted. Then, these chemicals were standardized by mapping to PubChem identifiers and Common Chemistry Registry Numbers (CASRN) (https://commonchemistry.cas.org/), resulting in 144 unique chemicals, of which 80 chemicals were from CTD and 66 chemicals from ECOTOX. Next, using PubChem identifiers, two-dimensional (2D) structures were obtained for these chemicals from PubChem (https://pubchem.ncbi.nlm.nih.gov/), along with their SMILES, InChI, and InChIKey representations. Subsequently, ClassyFire (Djoumbou Feunang et al., 2016) (http://classyfire.wishartlab.com) was used to classify the chemicals as organic or inorganic, resulting in the identification of 124 organic mercury compounds (Supplementary Table S1). Thereafter, SMILES strings corresponding to these chemicals were manually inspected, and those lacking dot separators (‘.’) were annotated as unfragmented.

### 2.2. Compilation and curation of AOPs within AOP-Wiki

An Adverse Outcome Pathway (AOP) represents a structured, sequential pathway detailing a stressor-induced toxicological mechanism (Ankley et al., 2010; Vinken et al., 2017). It begins with the interaction of a stressor and a biological receptor, commencing a Molecular Initiating Event (MIE) that culminates in an observed Adverse Outcome (AO) (Villeneuve et al., 2014a). The discrete nodes within this pathway are termed Key Events (KEs), while the relationships connecting them are designated as Key Event Relationships (KERs) (Ankley et al., 2010). Notably, KEs and KERs are supported by scientific evidence and organized across hierarchical biological organization levels (BOLs). AOP-Wiki (https://aopwiki.org) serves as a globally accessible, collaborative platform that facilitates the deposition and sharing of AOPs, along with their supporting evidence.

In this study, AOP-Wiki data was extracted from the downloadable XML file (released October 1, 2024), available on the ‘Project Downloads’ section of AOP-Wiki, utilizing an in-house Python parsing script. The resulting dataset comprised 487 AOPs, encompassing a total of 1485 KEs (Supplementary Table S2) and 2035 KERs (Supplementary Table S3). Extracted metadata included AOP identifiers, titles, associated KEs (including MIEs and AOs), KERs, linked stressors, the version of the OECD handbook utilized for development, AOP status as defined by OECD guidelines, biological applicability information (e.g., taxonomy, sex, life-stage), and corresponding weight of evidence (WoE) assessments. Furthermore, for each KE, data on title, identifier, level of biological organization, action name, object name, object identifiers, and associated process were extracted. Extracted KER data included upstream/downstream KEs, evidence supporting the biological plausibility of the relationship, adjacency information, and the degree of quantitative understanding attributed to the KER. Subsequently, 317 complete and connected AOPs were identified through a combination of systematic computational analysis and manual review (Sahoo et al., 2024a, 2024b), and were designated as curated AOPs. These 317 curated AOPs (Supplementary Table S4) comprised 1023 KEs (Supplementary Table S2) and 1573 KERs (Supplementary Table S3).

### 2.3. Compilation and curation of organic mercury-associated toxicity endpoints and protein targets

From published literature, CTD (Davis et al., 2023) (https://ctdbase.org/) has systematically curated species-specific associations for chemicals, phenotypes, diseases, and genes/proteins. In CTD, phenotypes are represented by Gene Ontology (GO) terms, and diseases are annotated with Medical Subject Headings (MeSH) identifiers. To identify fish-specific toxicity endpoints associated with organic mercury compounds, the fish-specific chemical-phenotype and chemical-disease associations were extracted from the January 2025 release of CTD, yielding 13 unique GO terms and 42 unique MeSH identifiers, respectively.

The ECOTOX database (Olker et al., 2022) provides a comprehensive, curated repository of ecotoxicological information for over 13,000 chemicals across more than 14,000 terrestrial and aquatic species. The complete dataset is available for download via the ‘Download ASCII Data’ option on the website. This dataset comprises information on test organisms, chemical identity, toxicity values, and observed toxicological effects, including biological effects (designated as ‘Effect’), corresponding measurement parameters (designated as ‘Measurement’), and effect trends (designated as ‘Trend’). The combination of ‘Measurement’ and ‘Trend’ provides a meaningful biological endpoint. In this study, the December 12, 2024, release of ECOTOX was downloaded, and subsequently, an in-house Python script was utilized to parse the fish-specific endpoint information from the ‘tests.txt’, ‘results.txt’, ‘chemicals.txt’, and ‘species.txt’ files. This process resulted in the identification of 238 unique toxicity endpoints for organic mercury compounds.

To identify proteins interacting with organic mercury compounds within CTD, fish-specific chemical-protein associations were filtered by restricting the ‘Geneform’ category to ‘protein’, resulting in the identification of 14 proteins. In parallel, the ECOTOX ‘Measurement’ list was manually screened for entries corresponding to organic mercury compounds in fish, resulting in the identification of 11 associated proteins. Overall, this systematic procedure resulted in the identification of 24 fish-specific protein targets for organic mercury compounds, which were further utilized for downstream analyses (Supplementary Table S5).

### 2.4. Molecular docking of organic mercury compounds to target proteins

In the AOP framework, the chemical-target protein interaction is indicative of the molecular initiating event (MIE), which then triggers the downstream events (Allen et al., 2014). Zebrafish (*Danio rerio*) offer significant advantages as a representative vertebrate model for aquatic organisms, owing to their fully sequenced genome, conserved molecular pathways, and demonstrated sensitivity to environmental toxicants (Horzmann and Freeman, 2018), making them well-suited for this investigation. It was observed that the identified target proteins for organic mercury compounds were from diverse fish species, limiting the scope for zebrafish proteins. Therefore, molecular docking was employed to predict binding affinities (in terms of predicted free energy of binding in kcal/mol) of organic mercury compounds with a range of zebrafish proteins, offering insights into previously uncharacterized toxicologically relevant molecular interactions. This subsection details the methodology employed for docking investigation, including data processing and analysis (Figure 1).

#### 2.4.1. Identification of zebrafish-specific target proteins

Given that the identified target proteins were not specific to zebrafish, an orthology assessment was undertaken using publicly available protein sequence resources. UniProt identifiers were first retrieved from the UniProt database (https://www.uniprot.org/), and these identifiers were subsequently queried against OrthoDB (https://www.orthodb.org/) to determine corresponding zebrafish orthologs, yielding 13 zebrafish orthologous sequences. To identify proteins not directly represented by UniProt identifiers in OrthoDB, a manual search of UniProt and UniParc (https://www.uniprot.org/uniparc) databases was conducted using protein names. Overall, 23 UniProt and one UniParc identifier were retrieved, representing zebrafish orthologs of the 24 identified target proteins (Supplementary Table S5).

#### 2.4.2. Compilation and curation of target protein structures

Molecular docking necessitates three-dimensional (3D) protein structures. First, the Protein Data Bank (PDB) (https://www.rcsb.org/) was queried to identify experimentally validated structures, resulting in the identification of a 3D structure for one protein. The AlphaFold Protein Structure Database (AlphaFold DB; https://alphafold.ebi.ac.uk/), a collaborative effort between DeepMind and EMBL-EBI, provides structural predictions for over 200 million UniProt sequences for which PDB entries are not available. Alphafold DB was queried using the UniProt identifiers, resulting in the identification of predicted 3D structures for 20 proteins.

For the remaining three proteins, *de novo* protein structure prediction was performed utilizing the ColabFold Jupyter notebooks (Mirdita et al., 2022). ColabFold provides a user-friendly implementation of AlphaFold2 (Jumper et al., 2021), a neural network-based model developed by DeepMind capable of generating structural models with accuracy approaching experimental resolution. The confidence in these predicted structures was assessed using the predicted Local Distance Difference Test (pLDDT) score (Vander Meersche et al., 2025), ranging from 0 to 100. Models exhibiting pLDDT scores above 90 were considered indicative of high local accuracy, while scores between 70 and 90 were deemed sufficient for structural modeling purposes (Jumper et al., 2021). Finally, to minimize contributions from intrinsically disordered regions, the terminal protein segments far from the protein core structures were truncated (Supplementary Table S5).

#### 2.4.3. Validation of obtained protein structures

The obtained protein structures were subjected to rigorous validation using Ramachandran plot analysis performed via the SAVES v6.0 server (https://saves.mbi.ucla.edu/), which incorporates PROCHECK (Laskowski et al., 1993) for stereochemical assessment. This analysis evaluates φ (phi) and ψ (psi) dihedral angles to assess the conformational plausibility of the protein backbone, as described by Laskowski *et al*. (Laskowski et al., 2018). All 24 3D structures demonstrated a significant portion of residues residing within favored and allowed regions of the Ramachandran plot (Supplementary Table S5), fulfilling criteria for structural soundness, and were therefore retained for subsequent analyses.

#### 2.4.4. Identification of ligand binding sites in the target proteins

To identify potential ligand binding sites in the target proteins, P2Rank (Jendele et al., 2019), a template-free and machine learning-based method, was employed. P2Rank was executed locally in batch mode to facilitate high-throughput pocket identification. The default prediction model was utilized, which incorporates both geometric and physicochemical properties of solvent-accessible surface residues. For each protein, ligandability scores were computed and clustered spatially to identify probable binding pockets. These pockets were then ranked according to their cumulative ligandability scores, with the top-ranked sites selected for subsequent molecular docking studies. A list of predicted pocket residues and associated scores is provided in Supplementary Table S6.

#### 2.4.5. Identification of reference biological ligands

BindingDB (https://www.bindingdb.org/) is a public database containing measured binding affinities for protein-ligand interactions primarily derived from the scientific literature. Therefore, BindingDB was queried with the 24 zebrafish proteins to identify biologically relevant reference ligands for docking studies. The resulting compounds for each target were manually reviewed, and ligands were selected based on the availability and strength of their affinity values. Compounds exhibiting high binding affinity (i.e., low K_i_, K_d_) were chosen as reference biological ligands for comparative docking studies and binding site validation (T. Liu et al., 2025).

Given the limited availability of zebrafish-specific ligand information, ligands derived from orthologous protein entries in other species were utilized. To justify this orthology-based protein identification process, a pairwise sequence alignment was conducted using EMBOSS Needle, which was accessed through the EMBL-EBI Job Dispatcher server (https://www.ebi.ac.uk/jdispatcher/), and a structural comparison was conducted using PyMOL version 2.5.0 (https://www.pymol.org/). Pairwise similarity scores obtained through EMBOSS Needle informed about the extent of sequence conservation (Supplementary Table S6). Furthermore, zebrafish protein structures predicted using AlphaFold were superimposed onto the crystal structures of their orthologs using PyMol to assess structural similarities based on root mean square deviation (RMSD) values. Low RMSD values (e.g., < 1.5 Å for most target proteins) indicated a high degree of structural conservation, supporting the translatability of binding pockets and validating the use of reference ligands identified from orthologous proteins in other species for docking experiments with zebrafish proteins (Supplementary Table S6). Through this extensive process, reference ligands were identified for 17 out of the 24 zebrafish proteins. These reference biological ligands, characterized by strong affinity and similarity to orthologous proteins from other species, were then used for docking investigation.

#### 2.4.6. Molecular docking of mercury compounds against the target proteins

Molecular docking of the organic mercury compounds against the target proteins in zebrafish was performed using AutoDock Vina version 1.2.3 (Eberhardt et al., 2021). To perform docking, the ligand and protein structures were converted from standard formats (e.g., SDF, PDB) to the PDBQT format using Open Babel version 3.1.0 (https://www.openbabel.org/), which also automated protonation, energy minimization, and file formatting steps.

For each target protein, docking grid boxes were defined using the top-ranked binding site coordinates obtained from P2Rank predictions. The search space center and grid box dimensions were assigned accordingly, and individual configuration files were automatically generated using custom shell scripts. Subsequently, protein-ligand docking was performed using AutoDock Vina by setting the exhaustiveness parameter to 12. The best docked poses (with the lowest binding energies) were obtained from AutoDock Vina results using the associated script *vina_split,* and a combined PDB structure of the protein-ligand complex corresponding to the best-docked pose was prepared using in-house Python scripts and pdb-tools (Rodrigues et al., 2018). Finally, predicted binding energies were compared between target protein-organic mercury compound pairs and target protein-reference ligand pairs. In this study, an organic mercury compound was considered to be binding with the target protein if its binding energy was equal to or lower than that of the reference biological ligand for the same protein.

### 2.5. Standardization of organic mercury-associated toxicity endpoints

Reported endpoints for chemicals can differ across toxicity databases, creating challenges for assessing organic mercury toxicity in fish and for comparing results between studies. To address this issue and harmonize endpoint descriptions, three controlled vocabularies or ontologies were utilized, namely, Key Events (KEs) from the AOP-Wiki, Gene Ontology (GO) terms (Ashburner et al., 2000; The Gene Ontology Consortium et al., 2023) (https://geneontology.org/), and Medical Subject Headings (MeSH) identifiers (https://www.nlm.nih.gov/mesh/meshhome.html). GO provides a standardized vocabulary for describing genes, gene products (e.g., proteins and non-coding RNAs), and their functions across species, encompassing molecular function, biological process, and cellular localization. MeSH identifiers are the National Library of Medicine’s controlled vocabulary thesaurus that provides descriptors used to index concepts in PubMed articles, organized within a hierarchical structure accessible via the MeSH Browser (https://meshb.nlm.nih.gov/).

The list of endpoints was compiled from previously curated information, including zebrafish proteins that successfully docked with at least one mercury compound in the docking investigation described in the previous subsection. To map these endpoints to the selected vocabularies, manual searches were conducted using the compiled KE list from AOP-Wiki, Amigo GO Browser (Carbon et al., 2009) (https://amigo.geneontology.org/), and MeSH Browser, respectively. It was observed that the KEs, GO terms, and MeSH identifiers, together, represented similar biological events, and therefore, were encapsulated as triplets representing those events. Thus, each mapped endpoint was assigned to at least one identifier among the triplet (KE, GO, MeSH) to ensure representation from each vocabulary. This facilitated the integration of diverse endpoint descriptions by linking them to common biological actions, acknowledging potential variations across terminologies.

In the AOP framework, KEs are organized by hierarchical BOLs, namely, Molecular, Cellular, Tissue, Organ, Individual, and Population, to convey mechanistic context (Villeneuve et al., 2014a, 2014b). As the GO terms and MeSH identifiers do not explicitly encode BOLs, they were manually inspected and assigned appropriate BOLs. Next, the compiled terms were assigned a node type such as MIE, AO, or KE based on the corresponding BOL, with terms in the Molecular level classified as MIEs, terms in Organ, Individual, or Population levels classified as AOs, and the remaining terms as KEs. This procedure yielded standardized organic mercury-associated toxicity endpoints with assigned identifiers, BOLs, and node types for subsequent analyses.

### 2.6. Construction of an organic mercury-associated toxicity network

Organizing toxicity information as a directed graph encoding sequential relationships among biological events can facilitate the identification of candidate pathways that can eventually be developed into computational AOPs (Ramšak et al., 2022). In this study, a novel integrative network-based approach was devised to construct such a toxicity network (Figure 2). This network comprises standardized organic mercury-associated toxicity endpoints as nodes and edges capturing relationships between different endpoints based on existing KER information and published evidence.

**Figure 2:**
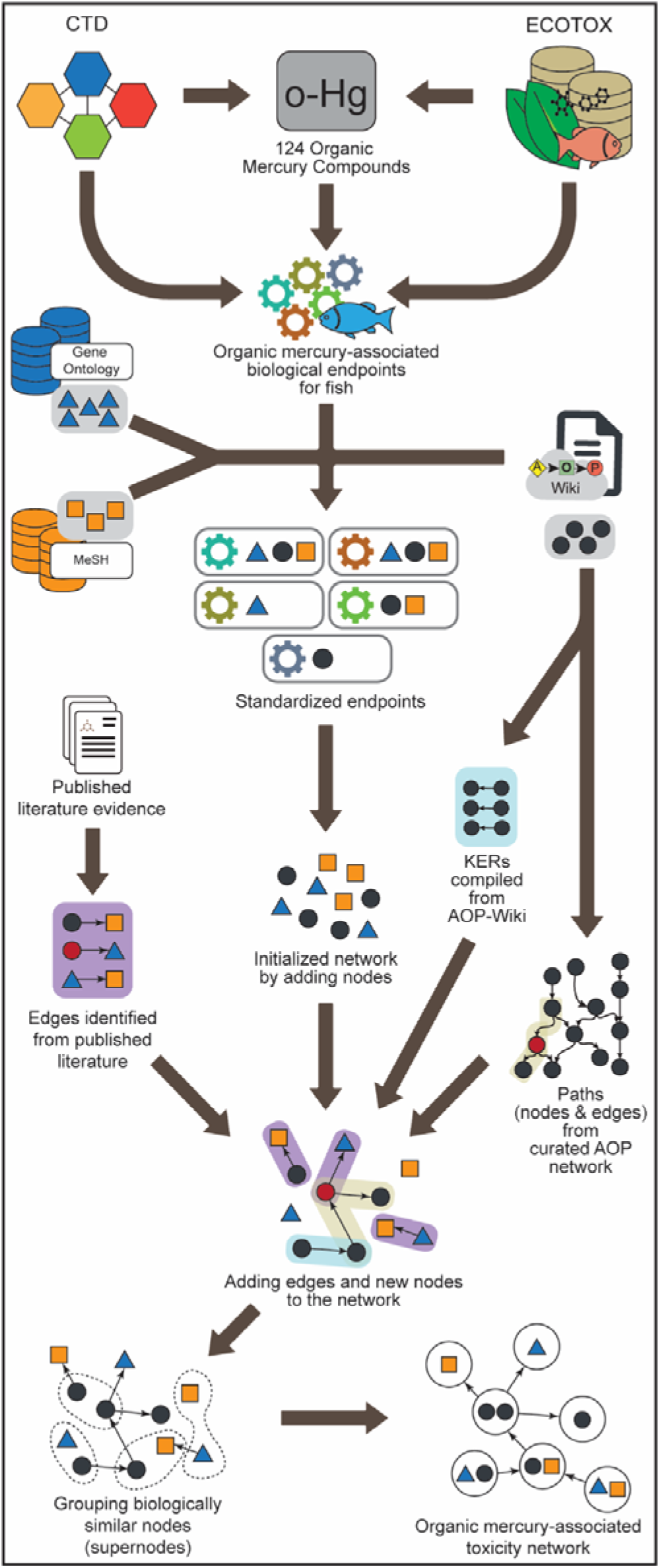
Integrated network-based workflow to construct an organic mercury-associated toxicity network.

#### 2.6.1. Identification of edges from existing KERs within AOP-Wiki

KERs are directed relationships that connect KEs within individual AOPs (Villeneuve et al., 2014a). To initialize edges in the organic mercury-associated toxicity network, KERs were added if their upstream and downstream KEs correspond to nodes already present within the toxicity network.

While KERs effectively capture the direct connections between KEs, relying solely on them may overlook the indirect links between different KEs. This limitation is addressed by constructing AOP networks, which reveal emergent paths extending beyond individual AOPs (Villeneuve et al., 2018). Accordingly, an AOP network was constructed using 317 curated AOPs, and thereafter, leveraged to identify novel paths between disconnected nodes in the toxicity network. First, KE pairs that remained disconnected following the KER integration were identified. Next, using the NetworkX (Hagberg et al., 2008) (https://networkx.org) module in Python, all possible shortest paths between each disconnected KE pair were determined within the curated AOP network. To ensure relevance to organic mercury toxicity, every KE on each path was screened for published evidence linking it to organic mercury exposure in ecological species. Only paths for which all constituent KEs were supported by such evidence were retained, with these KEs added to the set of nodes in the toxicity network, and the corresponding KERs incorporated into the edge list.

#### 2.6.2. Identification of edges from published evidence

To enable identification of links for GO terms and MeSH identifiers in the toxicity network that were not covered in KER information, this study relied on AOP-Helpfinder to extract evidence from published literature. AOP-Helpfinder (Carvaillo et al., 2019; Jaylet et al., 2025) is a text-mining tool that can extract event-event relationships using the PubMed database.

First, a comprehensive list of all the nodes in the toxicity network was generated and used as input for AOP-Helpfinder. The search parameters were configured with reduced search being set to ‘0%’, allowing a complete abstract search, and word proportion set to ‘standard method’, allowing partial matches between query terms and abstract content. Subsequently, the relationships identified by AOP-Helpfinder were manually verified, and only biologically plausible connections were retained. These verified relationships were then incorporated into the network’s edge list.

The identification of edges from existing KER information and published evidence facilitated the identification of both established and novel relationships between the nodes and significantly expanded the toxicity network’s connectivity.

#### 2.6.3. Grouping of biologically similar nodes in the toxicity network

While AOP-Wiki is the largest globally available repository of AOPs, the data within it is highly variable and heterogeneous, often resulting in overlapping and repetitive KE information (Mortensen et al., 2025; Wittwehr et al., 2025, 2024). Due to these redundancies and inconsistencies, AOP-Wiki does not fully comply with FAIR (Findable, Accessible, Interoperable, and Re-usable) data sharing principles (Wilkinson et al., 2016). As a consequence, functional redundancy was observed among KEs within the constructed toxicity network. Additionally, the network comprises GO terms and MeSH identifiers that are functionally similar to KEs, introducing further redundancy.

To address this challenge, a method was devised to group functionally similar nodes into a single entity. First, the KE, GO, and MeSH triplets associated with identified toxicity endpoints were considered. These triplets were then merged if they shared at least one identifier. These merged groups were then temporarily labeled and manually inspected to verify functional similarity. If a group represented a functionally similar action, it was designated as a ‘supernode’. Otherwise, the group was split into multiple subgroups, each potentially representing a distinct supernode based on functional similarity. This process provided a means of identifying and mitigating redundancy by consolidating nodes representing related functions.

Within the resulting supernode-level toxicity network, two supernodes were linked if their constituent nodes were connected via at least one edge from the original edge list, and these connections were termed ‘superedges’. Through this extensive process, a comprehensive toxicity network was constructed to provide the basis for understanding the candidate pathways. Subsequently, the candidate pathways were filtered, scored, and ranked for biological relevance and novelty, thereby facilitating the construction of ecologically relevant cAOPs for organic mercury-induced toxicity.

### 2.7. Construction of cAOPs for organic mercury-associated ecotoxicity

As the constructed toxicity network, comprising supernodes and superedges, contained multiple connected components, the largest connected component (LCC) was selected to maximize candidate pathway discovery. Next, supernodes containing at least one MIE or one AO were identified within this LCC. Subsequently, simple directed paths between all identified MIE-AO supernode pairs were enumerated using the NetworkX (Hagberg et al., 2008) (https://networkx.org) module in Python, resulting in the identification of potential candidate pathways.

In addition to MIE or AO classification, BOL information is necessary to ensure a logical progression from molecular events to adverse outcomes at higher levels (OECD, 2022), and is therefore essential to qualify candidate pathways as cAOPs. Since many supernodes contained mixed BOLs, evaluation was performed at the node level rather than assigning a single BOL to an entire supernode. Moreover, for each supernode pair connected in the candidate pathway, underlying nodes were examined for existing connections, and nodes without such links were removed (Figure 3). Novel links were then generated by substituting nodes with other members of the corresponding supernodes (Figure 3). Using these node-level links, all possible node-level paths were enumerated and treated as proxies for the supernode-level candidate pathways, while BOL annotations were maintained at every step (Figure 3).

**Figure 3:**
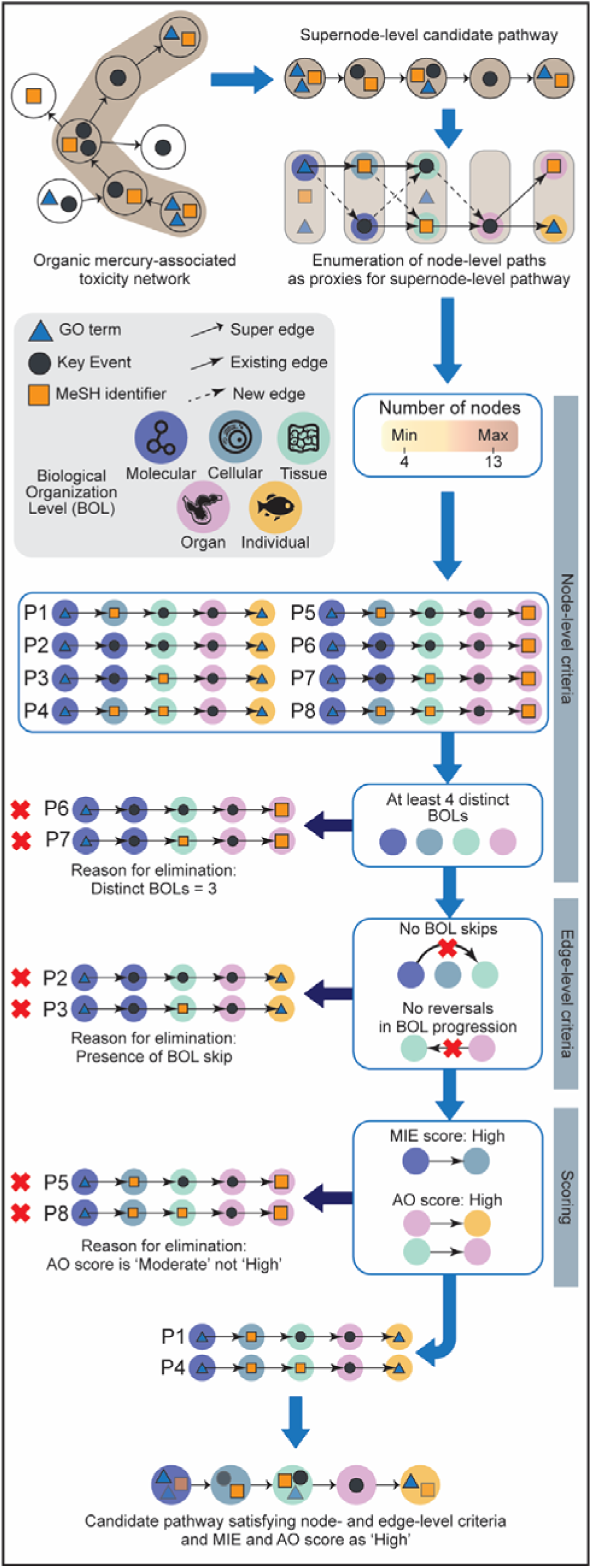
Filtration criteria for identification of candidate pathways for cAOPs.

Next, to ensure adherence to AOP definitions, these paths were subjected to various filtration criteria designed based on the guidelines presented in the AOP-Wiki Developers’ Handbook (OECD, 2022) (https://aopwiki.org/handbooks/5). The paths were first filtered based on the following node-level criteria:

● Minimum number of nodes in the path should be 4, which is the smallest possible standard AOP containing nodes with BOLs such as Molecular, Cellular, Tissue, and Organ.
● The nodes in the path should at least have 4 different BOL annotations, ensuring diversity consistent with standard AOP structure.
● Maximum number of nodes in the path should be 13, which is the largest number of nodes possible among the curated AOPs.

The edges in the resultant paths were reviewed to ensure that the downstream node was at the same level or higher than the BOL of the upstream node, and paths containing edges that moved down a level were removed. It was observed that in some paths, consecutive nodes were not in successive BOLs, indicating skips in the BOLs, and such paths were removed. Subsequently, to ensure BOL trajectory concordance, simple scores were devised based on the MIE and AO levels.

The MIE score is defined as follows:

● High, if the originating node is at Molecular level and the consecutive node is at Cellular level.
● Moderate, if the originating node is at Molecular level and the consecutive node is not at Cellular level, or the originating node is at Cellular level.
● Low, in other cases.

Similarly, the AO score is defined as follows:

● High, if the terminal node is at Population level, then the preceding nodes should be at Individual level, followed by at Organ level, or if the terminal node is at Individual level, then the preceding node should be at Organ level, or if the terminal node is at Organ level, then the preceding node should be at Tissue level.
● Moderate, if the terminal node is at Population, Individual, or Organ level, but its immediate predecessor node is not at Individual, Organ, or Tissue level, respectively.
● Low, in other cases.

The candidate pathways that satisfied all filtration criteria and achieved ‘High’ scores for both the MIE and AO were deemed consistent with AOP definitions and considered as candidate cAOPs. To assess novelty, a KER coverage score was computed for the underlying node-level paths, defined as the ratio of KERs overlapping with a curated AOP to the total number of KERs within the path. Since each candidate cAOP includes multiple node-level paths, the pathway’s coverage with respect to a given AOP was taken as the maximum coverage among its underlying node-level paths. To gauge the novelty of the candidate cAOP, a novelty score was then defined as 1 minus the maximum coverage across all curated AOPs. Subsequently, candidate cAOPs were ranked by this novelty score to identify the most novel pathways.

For candidate cAOPs to qualify as cAOPs, the biological plausibility of underlying nodes and edges was assessed using published evidence (Villeneuve et al., 2014a, 2014b; Yican Wang et al., 2024). Node plausibility was evaluated using published literature obtained using AOP-Helpfinder (Carvaillo et al., 2019; Jaylet et al., 2025) and Abstract Sifter (Baker et al., 2017). Edge plausibility was evaluated using published literature obtained through standardized PubMed queries (last searched on August 7, 2025), followed by a systematic literature review. Additionally, Bradford-Hill criteria (Becker et al., 2015; Collier et al., 2016) were utilized to understand the weight of evidence associated with each edge in these pathways. Through this extensive process, novel cAOPs were proposed for organic mercury-associated ecotoxicity in fish.

## 3. Results and Discussion

### 3.1. Molecular initiating events for organic mercury identified through docking-based investigation

This study aimed to construct computational Adverse Outcome Pathways (cAOPs) to explore organic mercury-induced toxicity in fish, and investigate their ecotoxicological risks, thereby supporting more informed risk assessment strategies for aquatic ecosystems. As a first step, molecular docking-based investigation was performed for 80 identified unfragmented organic mercury compounds (Methods; Supplementary Table S1) against 17 zebrafish proteins with reference ligand information (Methods; Supplementary Table S6), to gain insights into previously uncharacterized, toxicologically-relevant molecular interactions consistent with molecular initiating event (MIE) definition within the AOP framework (Allen et al., 2014; Villeneuve et al., 2014a).

It was observed that 16 zebrafish proteins exhibited successful docking with 44 organic mercury compounds (Methods; Supplementary Table S7). Among the docked proteins, Succinate dehydrogenase flavoprotein subunit (B2GQ13) and Heat shock protein 60 (Q803B0) had the highest number of successful ligand interactions, with 38 organic mercury compounds each, while ATP synthase subunit beta (A8WGC6) and Alkaline phosphatase (F1Q5B5) each has successful ligand interactions with only one organic mercury compound each (Supplementary Table S7).

Since AOP-Wiki (https://aopwiki.org/) is the largest repository of AOPs, which comprises data on MIEs among other biological events, these 16 binding-related events were checked against existing MIEs. It was observed that these identified events were not captured by the existing MIEs in AOP-Wiki and were therefore designated as novel MIEs and assigned a unique identifier for further analysis (Supplementary Table S8).

### 3.2. Organic mercury-associated toxicity network for fish

Toxicity networks constructed from chemical-specific information provide a basis for identifying candidate mechanistic paths that can be developed into cAOPs (Ramšak et al., 2022). In this study, a novel integrative network-based approach was devised to construct an organic mercury-associated toxicity network (Methods). First, 310 fish-specific toxicity endpoints were identified from CTD, ECOTOX, and molecular docking-based investigation (Methods). Thereafter, these endpoints were standardized by mapping to 143 KEs (including 16 novel MIEs), 102 GO terms, and 139 MeSH identifiers (Supplementary Table S8). The resulting 384 standardized endpoints were used as nodes to initialize the toxicity network.

Subsequently, the data from AOP-Wiki was relied upon to identify edges between these nodes within the toxicity network. First, 53 KERs were identified between 63 nodes and were added as edges to this toxicity network (Methods). Next, to expand the coverage, 860 paths were identified from the curated AOP network that connected 64 previously disconnected nodes (Methods). These 860 paths comprised 209 KERs and 137 KEs, of which 179 were novel KERs, and 73 were novel KEs with evidence of organic mercury-associated ecotoxicity (Supplementary Table S9). Hence, these 73 KEs and 179 KERs were added as nodes and edges into the toxicity network, respectively. Since AOP-Wiki data is limited to KER information, published evidence was next relied upon to identify links for GO terms and MeSH identifiers (Methods). This process resulted in the addition of 31 edges connecting 33 nodes in the toxicity network (Supplementary Table 10). Through this process, a toxicity network was constructed, comprising 457 nodes (Supplementary Table 11) and 263 edges (Supplementary Table 12).

Information in AOP-Wiki does not fully comply with FAIR (Findable, Accessible, Interoperable, and Re-usable) data sharing principles (Wilkinson et al., 2016) due to overlapping KE annotations (Mortensen et al., 2025; Wittwehr et al., 2025, 2024), and when combined with functionally similar GO terms and MeSH identifiers in our network, this leads to substantial redundancy. Therefore, these nodes were clustered into 197 functionally similar groups and were designated as supernodes (Methods). Subsequently, the edges were consolidated to capture the connectivity between supernodes and were designated as superedges (Methods). Altogether, through this extensive process, a fish-specific organic mercury-associated toxicity network was constructed, which comprised 197 supernodes (Supplementary Table 11) and 243 superedges (Supplementary Table 12).

### 3.3. Candidate cAOPs identified from the organic mercury-associated toxicity network

The constructed fish-specific organic mercury-associated toxicity network was examined to identify paths that can qualify as cAOPs. Since it was observed that the toxicity network consisted of disconnected components, the largest connected component (LCC), which comprised 123 supernodes and 235 superedges, was selected to maximize path discovery (Methods). As AOPs originate at an MIE and terminate at an adverse outcome (AO) (Villeneuve et al., 2014a), supernodes were first screened for these annotations. Within the LCC, it was observed that 30 supernodes contained at least one MIE node and 37 others contained at least one AO node, and were thus designated as MIE and AO supernodes, respectively.

In addition to MIE and AO annotations, AOPs include biological organization level (BOL) information to represent the progression from molecular events to adverse outcomes at higher levels (OECD, 2022). It was observed that many supernodes contained nodes with heterogeneous BOLs, making it difficult to characterize such supernodes. Therefore, a filtration procedure was devised to utilize the underlying node-level connections as proxies to understand the BOL progression at the supernode level (Methods). It was observed that 4865 paths, comprising 20 MIE supernodes and 14 AO supernodes, qualified the node-level and edge-level criteria (Methods). Among these, 973 paths with MIE and AO scores as ‘High’ (Methods), comprising 10 MIE supernodes and 11 AO supernodes, were considered further for cAOP construction (Supplementary Table S13).

In order to identify novel paths for cAOP construction, the obtained paths were compared with curated AOP information from AOP-Wiki (Methods). A novelty score was devised based on the edge overlaps with curated AOPs (Methods), and it was observed that four paths had the highest novelty score (Supplementary Table S13). Notably, these paths formed a network comprising 12 supernodes, including 4 supernodes that did not contain KEs, and 11 superedges, of which 5 represented novel links (Figure 4). It is important to note that AOPs, by definition, describe the simplest assembly of events, connecting a specific MIE to an AO, and are not intended to represent every possible perturbation (Villeneuve et al., 2014a). Thus, each MIE-AO connection can be considered the simplest path capturing biological events from a single MIE to a single AO, despite these paths forming a network. Therefore, these four paths were designated as candidate cAOPs (Figure 4).

**Figure 4:**
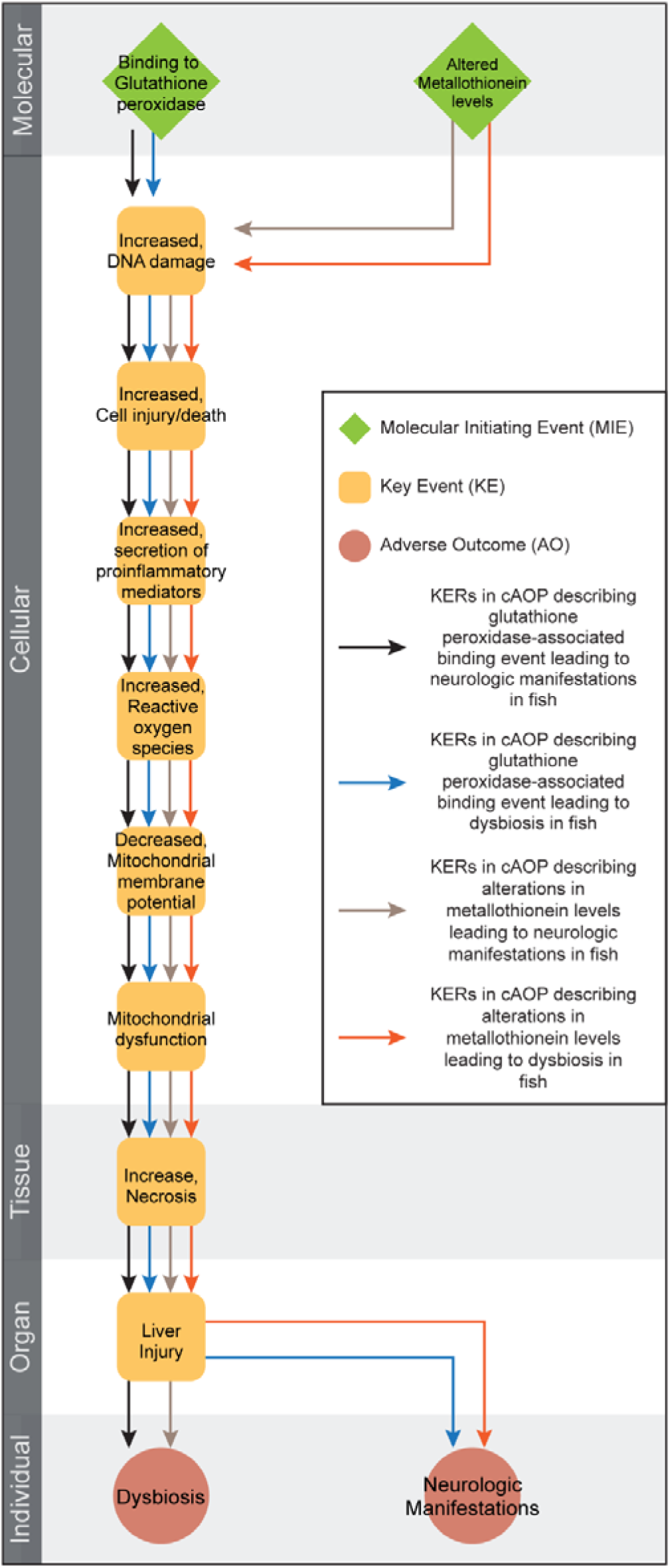
Directed network of four proposed cAOPs, comprising 12 nodes and 11 edges. Among the 12 nodes, 2 are categorized as MIEs (denoted as diamond), 2 are categorized as AOs (denoted as circle), and the remaining 8 are categorized as KEs (denoted as rounded square). The 11 edges are color-coded according to the specific cAOP to which they belong. In this figure, the 12 nodes are arranged vertically according to their level of biological organization.

### 3.4. Proposing ecotoxicologically-relevant cAOPs for fish

In order to assess the level of confidence in an AOP, it is necessary to understand the essentiality of the KEs and the biological plausibility of the KERs in the taxa of interest (Villeneuve et al., 2014a, 2014b; Yican Wang et al., 2024). In this study, published literature evidence was relied upon to assess the level of confidence of these four candidate cAOPs in fish (Figure 4; Supplementary Tables S14-S15).

#### 3.4.1. cAOP 1: Glutathione peroxidase-associated binding event leading to neurologic manifestations in fish

Glutathione peroxidase (GPx) is a key antioxidant enzyme that protects cells from oxidative stress (Formicki et al., 2025). In the docking-based investigation, 26 organic mercury compounds demonstrated strong binding energies toward zebrafish GPx (Supplementary Table S7). Notably, the compounds are bound to GPx with moderate yet comparable affinities relative to that of the reference ligand (Supplementary Table S7). For instance, Hydrargaphen (CID:16683105) showed a binding energy of −7.987 kcal/mol to GPx with hydrogen bond interactions to Tyr9, Glu162, and Glu164 residues (Figure 5a-b), while the reference ligand (BDB:51551531) showed a binding energy of −5.749 kcal/mol with hydrogen bond interactions to Tyr9 and Lys122 (Figure 5c-d). Moreover, in an earlier *in silico* study, Madabeni *et al*. explored the binding of methylmercury on cysteine and selanocysteine residues within GPx (Madabeni et al., 2021). These results suggest that organic mercury compounds could occupy or interact with the active sites of GPx. Further, published evidence supports that organic mercury compounds can inhibit GPx activity (Branco et al., 2012), suggesting binding to GPx as an MIE in this pathway.

**Figure 5:**
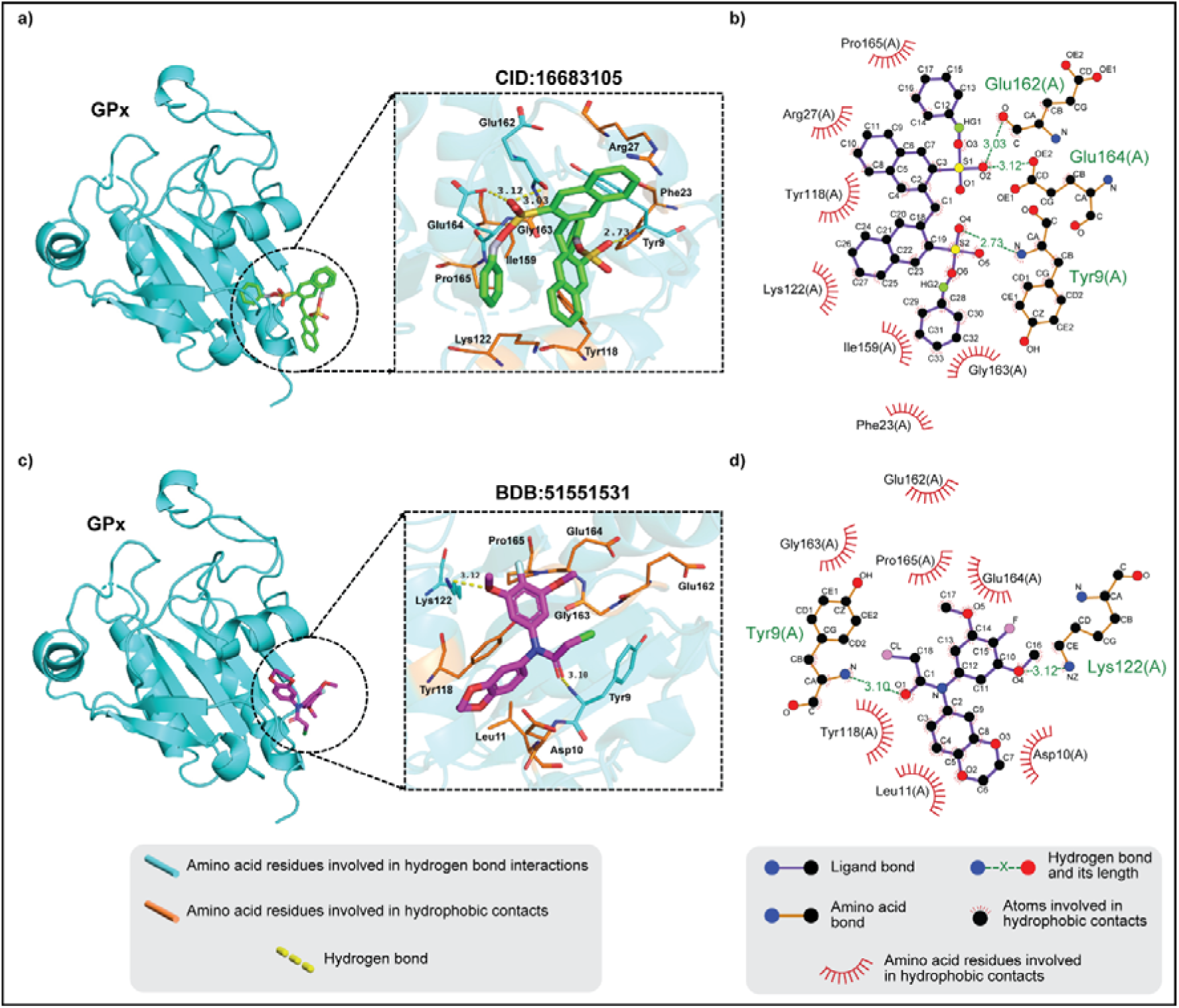
Cartoon representations and protein interaction diagrams of the glutathione peroxidase (GPx) protein in complex with Hydrargaphen (CID: 16683105) and reference ligand (BDB:51551531) in their best docked pose. **(a)** Cartoon representation of GPx-Hydrargaphen complex with inset highlighting ligand binding pocket, where the amino acid residues shown as blue sticks are involved in hydrogen bond interactions, and the amino acid residues shown as orange sticks are involved in hydrophobic contacts. **(b)** Protein interaction diagram of GPx-Hydrargaphen complex, where the interacting amino acid residues Tyr9, Glu162, and Glu164 are highlighted in green. **(c)** Cartoon representation of GPx-reference ligand complex with inset highlighting ligand binding pocket, where the amino acid residues shown as blue sticks are involved in hydrogen bond interactions, and the amino acid residues shown as orange sticks are involved in hydrophobic contacts. **(d)** Protein interaction diagram of GPx-reference ligand complex, where the interacting amino acid residues Tyr9 and Lys122 are highlighted in green. Here, the cartoon representations were generated using PyMol version 2.5.0 (https://www.pymol.org/), and the protein interaction diagrams were generated using LigPlot+ v1.4 (Laskowski and Swindells, 2011).

Inhibition of GPx has been observed to lead to DNA damage in fish (Ahmad et al., 2006; Ibrahim, 2015; Maharajan et al., 2018), initiating a cascade of cellular injury or death (Farag et al., 2006; Shaw et al., 2022) and increased secretion of pro-inflammatory mediators (Y. Chen et al., 2022; Ingerslev et al., 2010; Ogryzko et al., 2014). Further, it has been observed that elevated mediator levels in fish correlate with increased reactive oxygen species (ROS) levels (Baldissera et al., 2018; Cha et al., 2018; Souza et al., 2017), which subsequently disrupt mitochondrial membrane potential (J. Chen et al., 2022; L. Li et al., 2024; Sharaf et al., 2017) and lead to mitochondrial dysfunction (Datta et al., 2018; Qin et al., 2022; Saha et al., 2024). Histopathological analysis indicated that mitochondrial dysfunction can progress to hepatocyte necrosis (X. Liu et al., 2025) and eventually lead to liver injury in fish (Opute and Oboh, 2021; Suleiman et al., 2024). Subsequently, based on published evidence, correlations have been observed between liver injuries and neurologic manifestations such as reduced appetite, abnormal movement, lethargy, equilibrium loss, and erratic swimming in fish (Khan et al., 2025; Tahir et al., 2025; Topal et al., 2015; Ying Wang et al., 2024).

Furthermore, based on this published evidence (Supplementary Table S15), the weight of evidence for each KER in this cAOP was evaluated and presented in Table 1. Together, these findings support a cAOP describing the glutathione peroxidase–associated binding event leading to neurologic manifestations in fish.

**Table 1.**
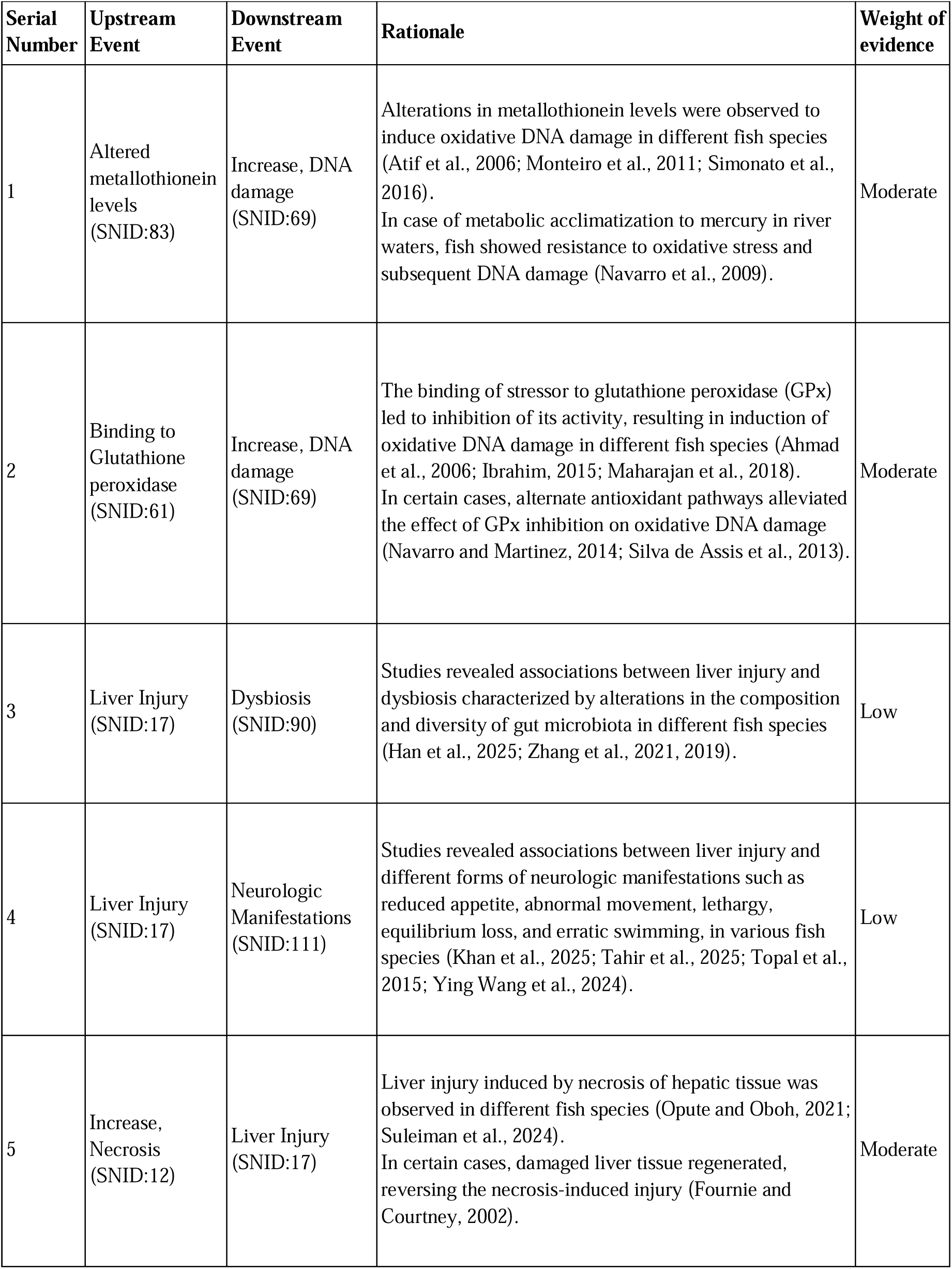
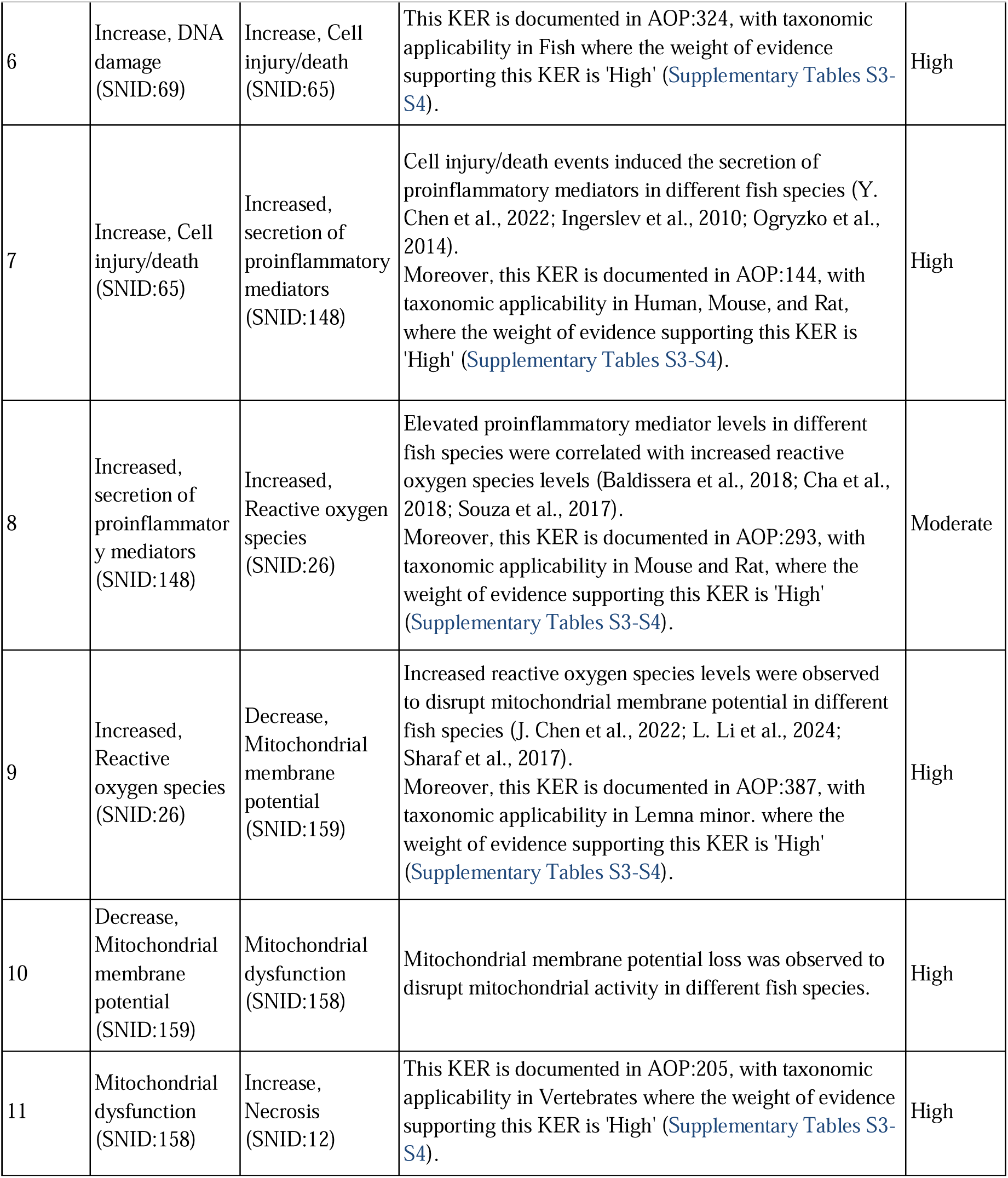
Weight of evidence assessment for the KERs present in the four proposed cAOPs.

#### 3.4.2. cAOP 2: Glutathione peroxidase-associated binding event leading to dysbiosis in fish

As presented in the previous cAOP, there is evidence of GPx binding causing oxidative stress that eventually leads to liver injury. This oxidative imbalance-mediated liver injury has been linked to dysbiosis in fish, which was characterized by significant alterations in composition and a decrease in diversity of gut microbiota with increased pathogenic bacteria (Han et al., 2025; Zhang et al., 2021, 2019). Collectively, these findings indicate that dysbiosis is an alternate adverse outcome that can potentially be triggered by GPx binding events. Table 1 presents the weight of evidence assessment based on this published data. Together, these findings support a cAOP describing the glutathione peroxidase–associated binding event leading to dysbiosis in fish.

#### 3.4.3. cAOP 3: Altered metallothionein levels leading to neurologic manifestations in fish

Metallothioneins (MTs) are small, cysteine-rich proteins widely induced by heavy metals in aquatic vertebrates (Vallee, 1991; Wang et al., 2014). They play a key role in detoxification by binding metal ions through thiol groups, thereby reducing the pool of redox-active metals that catalyze ROS formation (Vallee, 1991). When the tissue metal concentrations overwhelm MT levels, it can lead to accumulation of ROS (Sales Junior et al., 2024), and eventually DNA damage (Atif et al., 2006; Monteiro et al., 2011; Simonato et al., 2016). As presented in previous cAOPs, this can lead to oxidative imbalance-mediated neurologic manifestations in fish. Table 1 presents the weight of evidence assessment based on this published data. Together, these findings support a cAOP describing altered metallothionein levels leading to neurologic manifestations in fish.

#### 3.4.4. cAOP 4: Altered metallothionein levels leading to dysbiosis in fish

Based on the evidence supporting previous cAOPs, it was observed that altered metallothionein levels lead to dysbiosis through oxidative imbalance-induced liver injury.

Collectively, these findings indicate that dysbiosis is an alternate adverse outcome that can potentially be triggered by altered metallothionein levels. Table 1 presents the weight of evidence assessment based on this published data. Together, these findings support a cAOP describing altered metallothionein levels leading to dysbiosis in fish.

## 4. Conclusion

In this study, an integrative network-based framework was developed to accelerate the early-stage development of Adverse Outcome Pathways (AOPs) through the construction of computational AOPs (cAOPs), with a particular focus on elucidating the toxicity of organic mercury in fish. Specifically, 124 organic mercury compounds, corresponding fish-specific toxicity endpoints, and proteins were curated from publicly accessible toxicological databases. Next, novel molecular interactions were identified between organic mercury compounds and 16 zebrafish proteins. Thereafter, the toxicity endpoints and these novel molecular interactions were standardized and harmonized, and were designated as nodes to initialize the toxicity network. Subsequently, event relationships were compiled from AOP-Wiki and published literature, and were incorporated as edges to construct an organic mercury-associated toxicity network, comprising 197 nodes and 243 edges. Then, network-centric filtration criteria were designed based on AOP definitions and utilized to identify 973 biologically relevant pathways within the toxicity network. Finally, based on extensive evidence from published literature, four novel pathways that describe binding with glutathione peroxidase or alterations in metallothionein levels leading to neurologic manifestations or dysbiosis were proposed as cAOPs. Overall, this study demonstrates the utility of network-based methods that integrate existing ecotoxicological data to construct novel cAOPs.

While this study provides valuable insights, certain limitations should be noted regarding its wider applicability. The toxicological endpoint data were compiled exclusively from curated toxicological databases, which might not comprehensively capture all possible toxicological endpoints associated with the chemicals of interest. Molecular docking investigations were performed using zebrafish orthologous proteins that required structure predictions, and therefore, these predictions should be interpreted cautiously. This approach relies on existing AOP information within AOP-Wiki, which is subject to change over time as many AOPs are still under development and not yet endorsed. Moreover, the harmonization of toxicological endpoints required extensive manual review to resolve redundancies, thereby limiting the automation of this approach. The literature mining tools used in this study lacked organism-specific filters, necessitating additional manual effort to identify relevant studies.

Despite these limitations, this study proposes a novel computational network-based approach to support and streamline the AOP development process. A novel, comprehensive network elucidating organic mercury-specific toxicity mechanisms was constructed through the integration of ecotoxicological data from diverse resources. The harmonization of data from these resources enabled the identification of links that were previously missed due to node redundancies. The proposed criteria, derived from AOP definitions, provided a straightforward and effective method to assess the biological relevance of pathways within toxicity networks. Although organic mercury was used to construct the toxicity network and identify the cAOPs, the mechanisms were also observed with other stressors, showing that these cAOPs are stressor-agnostic.

In conclusion, this study presents a novel integrative network-based approach to construct cAOPs for fish, using organic mercury as a prototypical stressor, enabling targeted studies and potentially reducing the reliance on extensive animal testing. Furthermore, this methodology can be extended to broader range of contaminants and species, thereby enhancing New Approach Methodologies (NAMs) for toxicological risk assessment strategies.

## Data availability

The data associated with this study is contained in the article, or in the Supplementary Information.

## Acknowledgement

Areejit Samal would like to acknowledge funding from the Department of Atomic Energy (DAE), Government of India via Apex project to The Institute of Mathematical Sciences (IMSc) Chennai. This work was also undertaken as part of the project on ‘Marine Ecotoxicology and Ecological Risk Assessment’ (MEERA) Programme [MoES/OSMART/EFC/2021 dated 07.03.2022]. The authors are thankful to the Ministry of Earth Sciences (MoES), and the Director, National Centre for Coastal Research (NCCR), Government of India for the support and encouragement to carry out this work.

## CRediT author contribution statement

**Shreyes Rajan Madgaonkar:** Conceptualization, Data Curation, Formal Analysis, Methodology, Visualization, Writing; **Nikhil Chivukula:** Formal Analysis, Methodology, Visualization, Writing**; Vasavi Garisetti:** Data Curation, Formal Analysis, Methodology, Visualization, Writing**; Shambanagouda Rudragouda Marigoudar:** Conceptualization, Formal Analysis, Methodology, Writing**; Krishna Venkatarama Sharma:** Conceptualization, Formal Analysis, Methodology, Writing**; Areejit Samal:** Conceptualization, Supervision, Formal Analysis, Methodology, Writing.

## Supplementary Tables

**Table S1:** This table contains the list of 124 organic mercury compounds curated from publicly available chemical databases including CTD and ECOTOX. For each compound, this table provides the CAS Registry Number, PubChem chemical identifier (CID), database identifier(s), and chemical name. Further, for each compound, the table provides its structural representations in Canonical SMILES, InChI, and InChIKey format as reported in PubChem, and whether the chemical structure is unfragmented. The chemical taxonomy information (Kingdom, Superclass, and Class) was assigned based on the ClassyFire chemical classification system.

**Table S2**: This table contains information on the 1485 Key Events (KEs) compiled from the AOP-Wiki XML file released on October 1, 2024. For each KE, this table provides the corresponding information on KE identifier, KE title, biological organization level (BOL), associated action(s) (separated by ‘|’ symbol), object identifier(s) (separated by ‘|’ symbol) and name(s) (separated by ‘|’ symbol), and process identifier(s) (separated by ‘|’ symbol), source(s) (separated by ‘|’ symbol) and name(s) (separated by ‘|’ symbol) as provided in the XML file, and whether it is part of the curated list of 317 AOPs.

**Table S3**: This table contains information on Key Event Relationships (KERs) present in AOPs compiled from the AOP-Wiki XML file released on October 1, 2024. For each AOP, this table provides the AOP identifier, corresponding KER identifier, upstream KE identifier, downstream KE identifier, adjacency of KER, quantitative understanding of KER, weight of evidence of KER, and whether it is part of the curated list of 317 AOPs.

**Table S4**: This table contains the list of 317 curated adverse outcome pathways (AOPs) from AOP-Wiki. For each AOP, this table provides the corresponding information on AOP identifier, AOP title, Handbook Version that was followed while construction of the AOP,

Organisation for Economic Co-operation and Development (OECD) status, and taxonomic applicability of the AOP (separated by ‘|’ symbol).

**Table S5**: This table provides information on the 24 zebrafish orthologous protein targets of organic mercury compounds identified in this study. For each zebrafish orthologous protein, this table provides the protein identifier from UniProt or UniParc, the protein name, the curated target proteins along with their identifier, species in which it was identified and the database from where it was identified, the three-dimensional (3D) protein structure identifier along with the corresponding source from which it was identified, the sequence length of the protein, the average pLDDT score computed by AlphaFold along with the interpretation of the score, the residues that were truncated (if any), and the Ramachandran plot analysis information such as the residues in favoured regions (denoted by %), residues in additional allowed regions (denoted by %), residues in generously allowed regions (denoted by %), and residues in disallowed regions (denoted by %).

**Table S6**: This table contains the predicted binding pockets and associated reference ligands for the 24 zebrafish proteins considered for molecular docking. For each zebrafish protein, this table provides the protein identifier and name, predicted binding pockets along with the highest ligandability score obtained using P2Rank. Further, this table provides reference ligand information from BindingDB, such as the availability of information in the resource, associated orthologous protein (or Target Protein) name and organism in which it was identified, ligand identifier, ligand name, ligand structural descriptions (such as SMILES, InChI and InChiKey), and the corresponding reported inhibition constant (K_i_) (reported in ‘nM’), dissociation constant (K_d_) (reported in ‘nM’), and half maximal inhibitory concentration (IC_50_) (reported in ‘nM’). Furthermore, this table provides the three dimensional (3D) protein structure of the target protein from PDB or AlphaFoldDB, structural comparison between the zebrafish protein and target protein using the Root Mean Square Deviation (RMSD) (reported in Å), and the sequence comparison between the zebrafish protein and the target protein using the EMBOSS Needle scores such as identity (denoted as %), similarity (denoted as %), gaps (denoted as %), and overall score.

**Table S7**: This table contains the information on the obtained molecular docking-based binding energies for 16 zebrafish proteins. For each protein, this table provides the protein identifier and name, the corresponding docked ligand identifier, whether the docked ligand is considered a reference ligand, and docking-based binding energies (reported in kcal/mol).

**Table S8**: This table contains information on the organic mercury-associated toxicity endpoints from CTD, ECOTOX, and molecular docking. For each endpoint, this table provides the endpoint identifier and the corresponding source, the mapped Gene Ontology (GO) term and corresponding title, the mapped Medical Subject Heading (MeSH) identifier and corresponding title, the mapped KE identifier and corresponding title, and the assigned unique identifier (UID) and associated title for protein interactions with organic mercury compounds from molecular docking.

**Table S9**: This table contains information on the literature evidence of the 73 KEs that were added to support event-event associations between KEs that were mapped to organic mercury-associated toxicity endpoints. For each event, this table provides the corresponding KE identifier, KE title, biological organization level (BOL), organism in which organic mercury exposure is studied, study type, chemical that was tested, dosage information, abbreviated description of association with organic mercury exposure, and the corresponding reference.

**Table S10**: This table provides the event-event associations obtained from AOP-Helpfinder. For each event-event association, this table provides the identifier, title and reported actions for both upstream and downstream events, study type, organism in which the event-event association was observed, abbreviated description of the observed event-event association, and the corresponding reference.

**Table S11**: This table provides the node list for the organic mercury-associated toxicity network. For each node, this table provides their identifier, the type of the identifier, corresponding title, their biological organization level (BOL), and node types according to the AOP framework (MIE/KE/AO). Further, these nodes have been grouped based on event similarity and assigned to a unique supernode identifier (SNID).

**Table S12**: This table provides the edge list for the organic mercury-associated toxicity network. For each edge, this table provides the identifier and title for both the upstream and downstream events. Further, for each upstream and downstream node, this table provides the corresponding supernode identifier (SNID).

**Table S13**: This table contains information on candidate cAOPs obtained from the toxicity network. For each candidate cAOP, this table provides the corresponding path (where the nodes are separated by ‘|’ symbol), path length, corresponding MIE and AO, novelty score (rounded up to 6 decimals), and the maximum AOP coverage score with 317 curated AOPs (rounded up to 6 decimals).

**Table S14**: This table contains information on the literature evidence of the association with organic mercury for each of the 12 biological events present in the top-ranked candidate cAOPs. For each event, this table provides the corresponding event identifier (Supernode ID), event title (Supernode name), biological organization level (BOL), fish in which organic mercury exposure is studied, study type, chemical that was tested, dosage information, abbreviated description of association with organic mercury exposure, and the corresponding reference.

**Table S15**: This table contains information on the literature evidence of the biological plausibility of event-event relationships present in the top-ranked candidate cAOPs. For each event-event relationship, this table provides the corresponding upstream and downstream event identifier (Supernode ID) and event title (Supernode name), study type, fish in which the relationship was observed, chemical that was tested, dosage information, abbreviated description of the relationship and the corresponding reference, exceptions reported in the study, and brief description of the reported exception.

